# Impact of knowledge accumulation on pathway enrichment analysis

**DOI:** 10.1101/049288

**Authors:** Lina Wadi, Mona Meyer, Joel Weiser, Lincoln D. Stein, Jüri Reimand

## Abstract

Pathway-based interpretation of gene lists is a staple of genome analysis. It depends on frequently updated gene annotation databases. We analyzed the evolution of gene annotations over the past seven years and found that the vocabulary of pathways and processes has doubled. This strongly impacts practical analysis of genes: 80% of publications we surveyed in 2015 used outdated software that only captured 20% of pathway enrichments apparent in current annotations.

---

Pathway enrichment analysis is a common technique for interpreting candidate gene lists derived from high-throughput experiments^1^^-^^3^. It reveals the most characteristic biological processes and molecular pathways of candidate genes by combining statistical enrichment methods and prior knowledge of gene function from resources such as Gene Ontology^4^ (GO) and Reactome^5^. However, genomic and transcriptomic datasets have grown by several orders of magnitude since the turn of the century^6^,^7^ and the scientific literature has doubled between 2010 and 2015^8^. For example, GO is updated daily while new Reactome versions are released quarterly. Dozens of software tools have been written to interpret gene lists using information derived from GO and pathway databases, but only a few are regularly updated with the most current functional gene annotations. Many pathway enrichment tools have not been updated for years.

To investigate whether the age of pathway annotations adversely affects gene list interpretation, we analyzed the breadth and depth of functional information. We then reinterpreted cancer genes from recent experiments using gene annotation data with data snapshots taken at various times during the past seven years (2009-2016). First we surveyed the update times of 21 web-based pathway enrichment analysis tools^9^^-^^29^ (**Table 1**, **Supplementary Figure 1**). While several tools (g:Profiler^9^, Panther^12^, ToppGene^18^, GREAT^15^) were up to date with functional datasets revised within the past six months (February 2016), most publicly available tools were outdated by several years, and 12 (57%) tools had not been updated in the past five years (*e.g*. DAVID^19^ in January 2010). Nine tools had been updated within five years (*e.g*. FunSpec^29^, GeneCodis^23^) and eight tools had been updated within two years (*e.g*. Babelomics^14^, WebGestalt^11^). Surprisingly, the vast majority of surveyed publications from 2015 (84%) cited outdated tools that had not been updated within five years (**Figure 1A**, **Supplementary Table 1)**.

**Figure 1.**
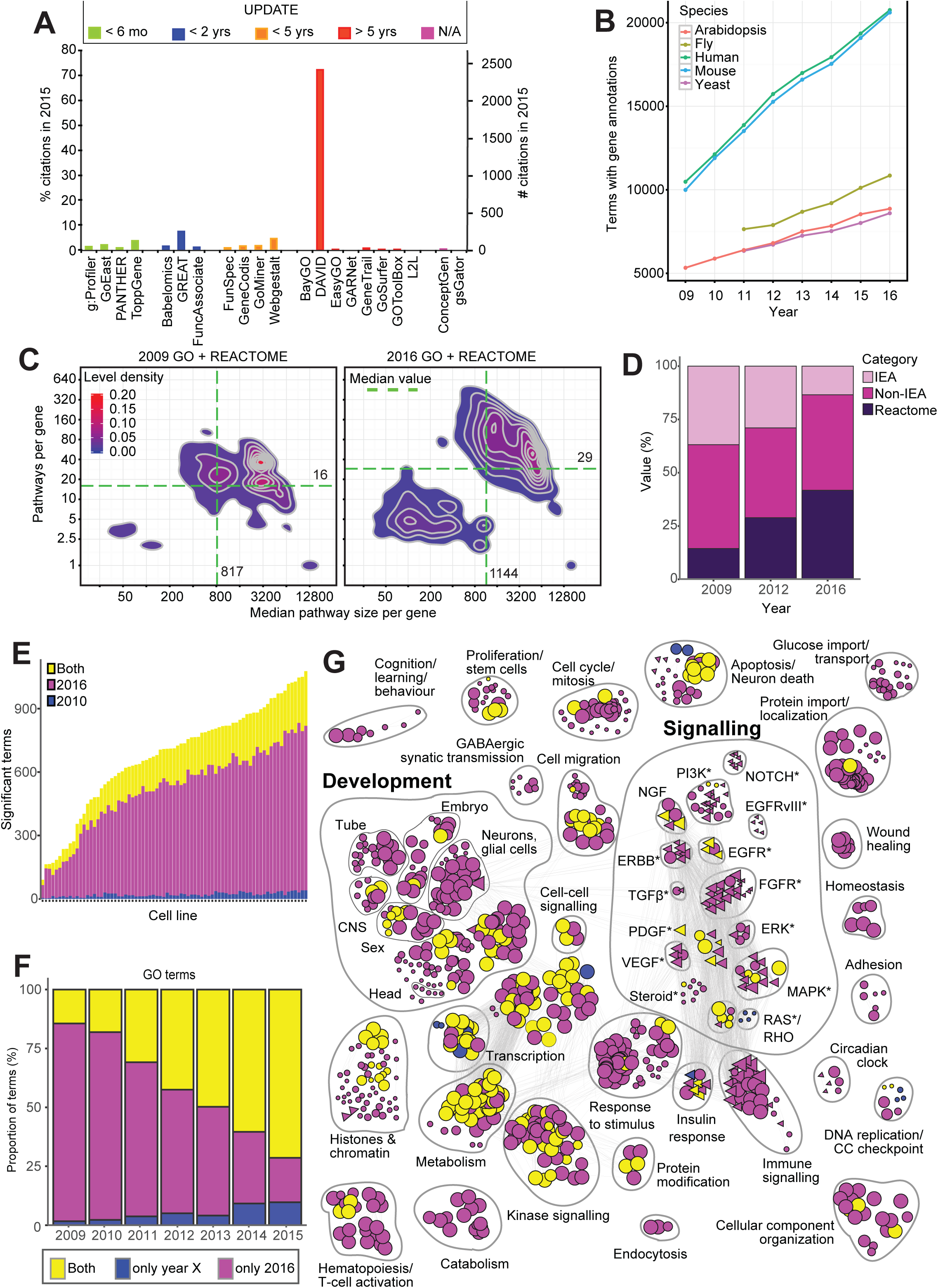
Pathway enrichment analysis relies on updated functional resources. **(A)** The majority of public pathway enrichment analysis software tools use outdated gene annotations, and the majority of surveyed papers published of 2015 use annotations that are more than 5 years old. **(B)** Annotated Gene Ontology (GO) terms per organism have grown rapidly in 2009-16 and more than doubled for human genes. **(C)** Two-dimensional density plots show accumulation of gene annotations during 2009-2016. Pathways per annotated gene (Y-axis) and median pathway sizes per annotated gene (X-axis) are shown. The median gene is shown with green dashed lines. **(D)** Quality of gene annotations is improving rapidly as manually curated Reactome annotations are becoming more frequent and fewer genes have only automatic GO annotations (Inferred from Electronic Annotation, IEA). **(E)** Gene annotations from 2010 miss 75% of enriched GO biological processes and Reactome pathways in essential breast cancer genes from recent shRNA screens. **(F)** Pathway enrichment analysis of frequently mutated glioblastoma (GBM) genes shows the proportion of results missed in outdated GO annotations. Each bar compares annotations of a given year to annotations of 2016. **(G)** Enrichment Map shows functional themes enriched in GBM genes that are missed in analysis of 2010 annotations. Nodes represent processes and pathways and edges connect nodes with many shared genes. Druggable pathways are indicated with asterisks.

**Table 1:**
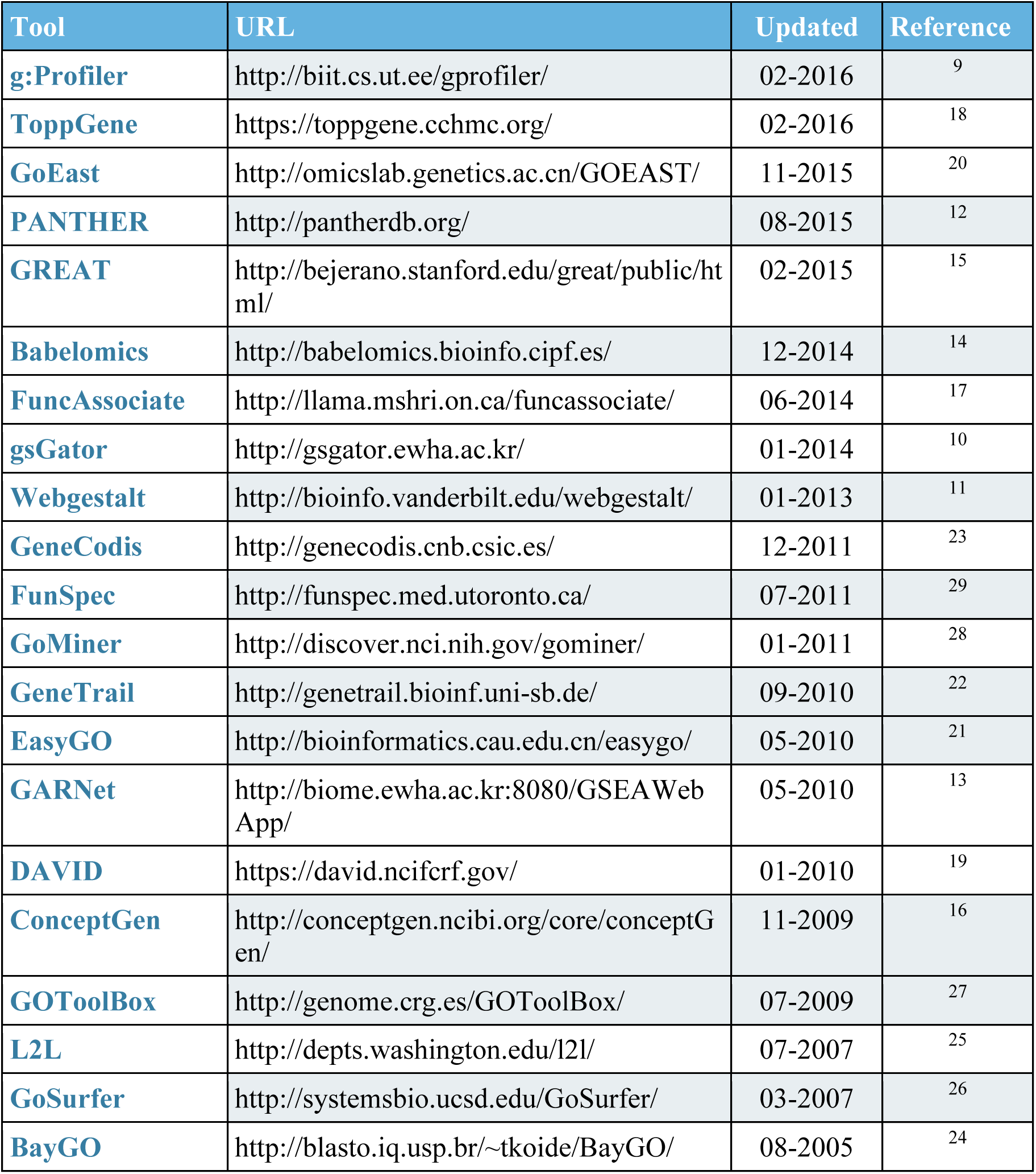
Web-based pathway enrichment analysis software tools with times of most recent updates.

We then asked how the use of outdated annotation databases would affect functional analysis of genes. We found consistent growth in the number and complexity of functional annotations over time across multiple species (2009-2016; **Figure 1B**). The number of biological processes in GO having human gene annotations has more than doubled between 2009 and 2016, from 6,509 to 14,735 with an average growth of 12.5% per year (**Supplementary Figure 2**). Similar growth was seen among cell components (936 to 1,728) and molecular functions (3,035 to 4,291). A typical pathway database, Reactome, had similar growth. Its human pathways nearly doubled over the same period from 880 to 1,746. GO annotations of mouse, fly, yeast, and *Arabidopsis thaliana* all grew at similar rates.

In addition to improvements in quantity, we found that the annotation databases have also improved in quality and complexity. The GO hierarchy has grown significantly deeper over time, as measured by the average path length between terms and root (7.59 to 8.06; *p*<10^−5^, permutation test; **Supplementary Figure 3**). The mean number of paths to root per term has tripled (47 to 160; *p*<10^−5^; **Supplementary Figure 4**). The former metric shows greater detail of our biological vocabulary, while the latter shows increasing interconnectedness of concepts. These changes directly affect gene list interpretation, as genes in GO terms are automatically propagated to parent terms.

Analysis of gene-term associations showed rapid growth in the knowledge of individual genes (**Figure 1C**). The median human gene in 2016 is associated with 29 processes and pathways compared 16 in 2009 (*p<*10^−5^). The median functional gene set has also grown from 817 to 1,144 genes (*p<*10^−5^). Earlier general terms included thousands of genes, while recent annotations cover a wide spectrum of broad and specific pathways and processes. In particular, the manually-curated Reactome resource has grown from 49 to 150 genes per median pathway (**Supplementary Figure 5**). The increase in GO annotations is also mirrored in model organisms (**Supplementary Figure 6**).

High-confidence experimental annotations are becoming more common, while the fraction of poorly annotated human genes is decreasing (**Figure 1D**). In 2016, 42% of genes have at least one Reactome annotation (versus 15% in 2009), while 14% of genes have only low-confidence electronic annotations (versus 37% in 2009). The first category is based on manual literature curation, while the latter only includes IEA annotations in GO (Inferred from Electronic Annotation). The ‘dark matter’ of the genome also contributes to the quality and quantity of gene annotations. In 2009, one of eight human protein-coding genes (12.4%) had no annotations in GO or Reactome, compared to 4.9% of genes in 2016 (**Supplementary Figure 7**). Changes in gene symbols also contribute to loss of information. Earlier software would miss one of eight genes due to changing gene nomenclature. Using standard gene symbols from 2015, we found that genes were missed due to nomenclature changes 12.2% of the time when using symbols from 2009, compared to just 1.1% of gene symbols in 2014 (**Supplementary Figure 8**). Collectively, these data show that gene annotations have become substantially broader, more specific and higher-quality over the last seven years, covering more protein-coding genes and reducing the number of un-annotated genes.

Given this study improvement, we evaluated the impact of annotation evolution on gene list interpretation. First we analyzed essential genes of 77 breast cancer cell lines derived from recent shRNA screens^30^. Strikingly, 74% of enriched terms of 2016 were missed when testing the top 500 essential genes from each cell line with annotations from 2010 (191 pathways per median cell line in 2010 *vs*. 695 in 2016, **Figure 1E**, **Supplementary Figure 9**). We also confirmed this finding using the top-100 genes (**Supplementary Figure 10**) and by repeating the analysis separately for GO and Reactome (**Supplementary Figure 11**).

To confirm our observations in a high-confidence dataset, we studied 75 significantly mutated driver genes in the glioblastoma (GBM) form of brain cancer, taken from a recent pan-cancer analysis of 6,800 tumors^31^,^32^ that includes both known cancer drivers (*EGFR*, *PIK3CA*, *PIK3R1*, *PTEN*, *TP53*, *NF1*, *RB1*)^33^ and less well-known candidates. A standard enrichment analysis, performed using Fisher’s exact test and outdated annotations, missed many pathways detected with 2016 data (**Figure 1F**, **Supplementary Figure 12, Online Methods**). In particular, the 2010 era annotations (which the DAVID software uses) only capture ~20% of the 2016 results, including 172/827 GO biological processes and 16/128 Reactome pathways. The 13 general processes exclusively seen in 2010, which include large groupings such as transcription and apoptosis, were restructured in later databases.

Some enriched pathways and processes are also missed in relatively recent gene annotations (**Figure 1F**). In the enrichment analysis performed using 2015 data, 89/743 (12%) of GO terms and 29/116 (25%) Reactome pathways were missed compared to 2016. Some missing GO terms (8%) are present in the set of annotations from 2016, while most are not significant at FDR *p*<0.05. However ~40% of the insignificant processes are seen at less stringent cutoffs (FDR *p*<0.1) (**Supplementary Figure 13**). As pathways grow over time, enrichment signals from gene lists are diluted. Thus researchers may not be able to replicate all pathways when repeating analysis using newer gene annotations.

A detailed summary of GBM pathways represented as an Enrichment Map^34^ shows major functional themes missed in 2010 annotations (FDR *p*<0.05, **Figure 1G**; **Supplementary Tables 2-3**). Certain themes neurotransmitter signaling (*n*=6, FDR *p*=0.0013), circadian clock (*n*=8, FDR *p*=1.5x10^−4^) and glucose signaling (*n*=7, FDR *p*=0.0016) are only highlighted in current analysis. These processes are expected from brain cancer genes ^35^,^36^, for example enhanced glucose uptake of brain tumor initiating cells helps these overcome nutrient deprivation^37^. In particular, immune response (*n*=29, FDR *p*=5.2x10^−5^) and related processes are only apparent in annotations from 2016 and emphasize emerging opportunities in cancer immunotherapy^45^. Other themes underline increased specificity of neuronal context: while apoptosis is highlighted in both analyses, neuronal apoptosis only comes up in newer data (*n*=7 genes, FDR *p*=0.018). The difference between annotations of 2010 and 2016 is even stronger when excluding GO IEA annotations, as 96.5% (603/625) of high-confidence pathways and processes are missed when older data are used (**Supplementary Figure 14**).

The analysis with current annotation data also highlights signaling pathways relevant to GBM biology and therapy development^38^, ^39^, ^33^ such as Notch (*n*=5, FDR *p*=0.0019), TGF-β (*n*=5, FDR *p*=0.027), and fibroblast growth factor (*n*=12, FDR *p*=1.13x10^−6^) (**Figure 1G**). For example, the Notch pathway is targetable with γ-secretase inhibitors (R04929097) that are currently in phase I and II glioma trials^38^. The TGF-β pathway can be inhibited via the ligand (Trabedersen) or the receptor (Galunisertib)^40^. Current gene annotations also highlight the EGFRvIII^41^,^42^ signaling pathway (*n*=5, FDR *p*=1.07x10^−5^) that is among the most common alterations in GBM. It involves deletion of exons 2-7 of the *EGFR* gene that drives tumor progression and correlates with poor prognosis^43^. While EGFR inhibitors have been unsuccessful in GBM treatment to date, the recently developed Rindopepimut vaccine targets EGFRvIII and has entered clinical trial^44^. These targetable pathways would not have been identified using outdated annotation information.

The growth in the quantity and completeness of functional annotations has a crucial impact on practical analysis of high-throughput data in current literature. Out of the 21 tools we studied, the most popular software was DAVID, used in 2,500 publications in 2015 and representing 71.4% of the citations of software tools we reviewed. Our analysis shows that at least 74% of pathway enrichment hits are missed when analyzing gene annotations from 2010, including ~12% of dark-matter genes lost due to lack of annotations, ~12% of genes lost due to outdated or absent gene symbols, and at least 50% of results missed due to enhancements in the catalog of biological pathways and processes. The implication is that thousands of recent studies have severely underestimated the functional significance of their gene lists because of outdated software and gene annotations. We now have the opportunity to discover new valuable information and outline experimental hypotheses by carefully re-analyzing existing datasets. The bioinformatics community needs to prioritize the timeliness of gene annotations and data reproducibility. At least semiannual software updates are required as genome databases and biomedical ontologies receive major updates several times a year. To ensure reproducibility of earlier analyses, software tools need to provide access to historical versions of gene annotations. As an example of recommended practice, our g:Profiler web server (http://biit.cs.ut.ee/gprofiler), which includes gene annotations for human and more than two hundred other organisms, is synchronized with the Ensembl database every few months and maintains an archive of earlier releases dating back to 2011.

Researchers and reviewers need to pay attention to timeliness of data, and software tools need to clearly indicate the time of the most recent update. Reliable and up-to-date software tools allow researchers to make the best use of current knowledge of gene function and best interrogate experimental data for making scientific discoveries.

## Online methods

### Ontologies and pathways

Functional terminology of biological processes, molecular functions, and cell components was retrieved from the Gene Ontology^4^ website and comprised January releases of each year (2009-2016). Gene annotations were derived from the Gene Ontology Annotation (UniProt-GOA) database^46^. Molecular pathways from the Reactome^5^ database were retrieved from archives and included December releases of previous years. Genes were annotated to GO terms as well as parent and ancestor terms via all possible paths. Obsolete terms and negative relationships in GO were removed. We filtered human genes with non-public status and analyzed protein-coding genes of matched versions of the NCBI Consensus Coding Sequence Database^47^ (CCDS).

### Analysis of gene annotations

Pathway databases were analyzed for growth in total number of pathway terms separately for the three main ontologies in GO (biological processes, molecular functions, cell components) for each year (2009-2016). The same analysis was repeated for human Reactome pathways and GO annotations for model organisms (mouse, *Arabidopsis thaliana*, fly, yeast). We counted GO terms and Reactome pathways with at least one annotated gene of the studied species. Path lengths and numbers from terms to roots were computed with custom scripts. Human annotations of GO terms and Reactome pathways contained high-confidence protein-coding genes from the nearest previous release of the CCDS database release (e.g. 2015 release for 2016 annotations) and only included genes with public status. Density of human gene annotations was assessed with two-dimensional density plots. For each gene, number of associated processes and pathways (Y-axis) and median size of corresponding gene sets per gene (Y-axis) were shown. The density plots include non-annotated genes (i.e. “dark matter”) for density estimation but are not shown. Dark matter genes were selected as protein-coding genes of the corresponding CCDS release that had no annotations in GO or Reactome, or only had top-level GO annotations (one or more of biological_protess, cell_component, molecular_function). GO biological processes per gene were also estimated for model organisms (mouse, *A. thaliana*, fly, yeast) without filtering of CCDS genes and dark matter. The proportion of missing gene symbols was estimated from earlier CCDS releases relative to the most recent CCDS release of 2015. Quality of human gene annotations was assessed in three mutually exclusive categories - genes with at least one Reactome annotation, genes with at least one nonelectronic (non-IEA) annotation in GO, and genes with only IEA (Inferred from Electronic Annotation) annotations in GO.

### Pathway Enrichment Analysis

Pathway enrichment analysis was conducted on GO biological processes and Reactome pathways using Fisher’s exact tests. Multiple testing was conducted separately for GO and Reactome terms using the Benjamini-Hochberg False Discovery Rate (FDR) procedure. Terms with FDR *p*<0.05 were considered significant. Enrichment analysis of GO and Reactome terms conservatively comprised separate background gene sets that included all the genes with at least one gene annotation GO Biological Process and Reactome, respectively. We chose this general enrichment strategy, as direct comparison of tools would be confounded by differences in underlying methods.

Two sets of enrichment analyses were conducted on cancer gene lists using gene annotations from 2016 (corresponding to g:Profiler) and 2010 (corresponding to DAVID). First we analyzed essential breast cancer genes from recent shRNA screens of 77 cell lines^30^. We separately analyzed top-100 and top-500 lists of genes according to per-gene zGARP scores provided by the study. We counted shared, outdated-only, and recent-only gene annotations enriched in the analyses (FDR *p*<0.05) and matched these using GO and Reactome term identifiers. The most common terms only found in the up-to-date analysis were visualized with the WordCloud R package. To simulate practical analysis, we did not manually convert outdated gene symbols in breast cancer analysis. Second, the same comparison pipeline was repeated for a smaller set of 75 glioblastoma driver genes found as frequently mutated in pan-cancer analysis^31^,^32^ derived from the IntOGen database^48^. We manually mapped outdated gene symbols in the GBM analysis to create a more conservative analysis scenario. We compared enriched annotations across the years 2009-2015 relative to 2016 and counted common and distinct terms as above. We also visualized the pathway and process enrichments of 2010 and 2016 using the Enrichment Map^34^ app of Cytoscape^49^. The Enrichment Map analysis covered pathways with at least four genes. Our observations were also confirmed when all pathways were included. Functional themes and signaling pathways were curated manually.

## References

1 Creixell, P. et al. Pathway and network analysis of cancer genomes. Nat Methods 12, 615–621, doi:10.1038/nmeth.3440 (2015).

2 Huang da, W., Sherman, B. T. & Lempicki, R. A. Bioinformatics enrichment tools: paths toward the comprehensive functional analysis of large gene lists. Nucleic Acids Res 37, 1–13, doi:10.1093/nar/gkn923 (2009).

3 Khatri, P., Sirota, M. & Butte, A. J. Ten years of pathway analysis: current approaches and outstanding challenges. PLoS Comput Biol 8, e1002375, doi:10.1371/journal.pcbi.1002375 (2012).

4 Ashburner, M. et al. Gene ontology: tool for the unification of biology. The Gene Ontology Consortium. Nature genetics 25, 25–29, doi:10.1038/75556 (2000).

5 Croft, D. et al. The Reactome pathway knowledgebase. Nucleic Acids Res 42, D472–477, doi:10.1093/nar/gkt1102 (2014).

6 The 1000 Genomes Project Consortium. A map of human genome variation from population-scale sequencing. Nature 467, 1061–1073 (2010).

7 Kolesnikov, N. et al. ArrayExpress updateߞsimplifying data submissions. Nucleic acids research 43, D1113–1116, doi:10.1093/nar/gku1057 (2015).

8 Powell, K. Does it take too long to publish research? Nature 530, 148–151, doi:10.1038/530148a (2016).

9 Reimand, J., Kull, M., Peterson, H., Hansen, J. & Vilo, J. g:Profilerߞa web-based toolset for functional profiling of gene lists from large-scale experiments. Nucleic acids research 35, W193–200, doi:10.1093/nar/gkm226 (2007).

10 Kang, H. et al. gsGator: an integrated web platform for cross-species gene set analysis. BMC Bioinformatics 15, 13, doi:10.1186/1471-2105-15-13 (2014).

11 Wang, J., Duncan, D., Shi, Z. & Zhang, B. WEB-based GEne SeT AnaLysis Toolkit (WebGestalt): update 2013. Nucleic Acids Res 41, W77–83, doi:10.1093/nar/gkt439 (2013).

12 Mi, H., Muruganujan, A. & Thomas, P. D. PANTHER in 2013: modeling the evolution of gene function, and other gene attributes, in the context of phylogenetic trees. Nucleic acids research 41, D377–386, doi:10.1093/nar/gks1118 (2013).

13 Rho, K. et al. GARNETߞgene set analysis with exploration of annotation relations. BMC Bioinformatics 12 Suppl 1, S25, doi:10.1186/1471-2105-12-S1-S25 (2011).

14 Medina, I. et al. Babelomics: an integrative platform for the analysis of transcriptomics, proteomics and genomic data with advanced functional profiling. Nucleic Acids Res 38, W210–213, doi:10.1093/nar/gkq388 (2010).

15 McLean, C. Y. et al. GREAT improves functional interpretation of cis-regulatory regions. Nature biotechnology 28, 495–501, doi:10.1038/nbt.1630 (2010).

16 Sartor, M. A. et al. ConceptGen: a gene set enrichment and gene set relation mapping tool. Bioinformatics 26, 456–463, doi:10.1093/bioinformatics/btp683 (2010).

17 Berriz, G. F., Beaver, J. E., Cenik, C., Tasan, M. & Roth, F. P. Next generation software for functional trend analysis. Bioinformatics 25, 3043–3044, doi:10.1093/bioinformatics/btp498 (2009).

18 Chen, J., Bardes, E. E., Aronow, B. J. & Jegga, A. G. ToppGene Suite for gene list enrichment analysis and candidate gene prioritization. Nucleic acids research 37, W305–311, doi:10.1093/nar/gkp427 (2009).

19 Huang da, W., Sherman, B. T. & Lempicki, R. A. Systematic and integrative analysis of large gene lists using DAVID bioinformatics resources. Nature protocols 4, 44–57, doi:10.1038/nprot.2008.211 (2009).

20 Zheng, Q. & Wang, X. J. GOEAST: a web-based software toolkit for Gene Ontology enrichment analysis. Nucleic Acids Res 36, W358–363, doi:10.1093/nar/gkn276 (2008).

21 Zhou, X. & Su, Z. EasyGO: Gene Ontology-based annotation and functional enrichment analysis tool for agronomical species. BMC Genomics 8, 246, doi:10.1186/1471-2164-8-246 (2007).

22 Backes, C. et al. GeneTrailߞadvanced gene set enrichment analysis. Nucleic Acids Res 35, W186–192, doi:10.1093/nar/gkm323 (2007).

23 Carmona-Saez, P., Chagoyen, M., Tirado, F., Carazo, J. M. & Pascual-Montano, A. GENECODIS: a web-based tool for finding significant concurrent annotations in gene lists. Genome Biol 8, R3, doi:10.1186/gb-2007-8-1-r3 (2007).

24 Vencio, R. Z., Koide, T., Gomes, S. L. & Pereira, C. A. BayGO: Bayesian analysis of ontology term enrichment in microarray data. BMC Bioinformatics 7, 86, doi:10.1186/1471-2105-7-86 (2006).

25 Newman, J. C. & Weiner, A. M. L2L: a simple tool for discovering the hidden significance in microarray expression data. Genome Biol 6, R81, doi:10.1186/gb-2005-6-9-r81 (2005).

26 Zhong, S. et al. GoSurfer: a graphical interactive tool for comparative analysis of large gene sets in Gene Ontology space. Appl Bioinformatics 3, 261–264 (2004).

27 Martin, D. et al. GOToolBox: functional analysis of gene datasets based on Gene Ontology. Genome Biol 5, R101, doi:10.1186/gb-2004-5-12-r101 (2004).

28 Zeeberg, B. R. et al. GoMiner: a resource for biological interpretation of genomic and proteomic data. Genome Biol 4, R28 (2003).

29 Robinson, M. D., Grigull, J., Mohammad, N. & Hughes, T. R. FunSpec: a web-based cluster interpreter for yeast. BMC Bioinformatics 3, 35 (2002).

30 Marcotte, R. et al. Functional Genomic Landscape of Human Breast Cancer Drivers, Vulnerabilities, and Resistance. Cell 164, 293–309, doi:10.1016/j.cell.2015.11.062 (2016).

31 Tamborero, D. et al. Comprehensive identification of mutational cancer driver genes across 12 tumor types. Sci Rep 3, 2650, doi:10.1038/srep02650 (2013).

32 Rubio-Perez, C. et al. In silico prescription of anticancer drugs to cohorts of 28 tumor types reveals targeting opportunities. Cancer Cell 27, 382–396, doi:10.1016/j.ccell.2015.02.007 (2015).

33 Brennan, C. W. et al. The somatic genomic landscape of glioblastoma. Cell 155, 462–477, doi:10.1016/j.cell.2013.09.034 (2013).

34 Merico, D., Isserlin, R., Stueker, O., Emili, A. & Bader, G. D. Enrichment map: a network-based method for gene-set enrichment visualization and interpretation. PloS one 5, e13984, doi:10.1371/journal.pone.0013984 (2010).

35 D’Urso, P. I. et al. miR-155 is up-regulated in primary and secondary glioblastoma and promotes tumour growth by inhibiting GABA receptors. International journal of oncology 41, 228–234, doi:10.3892/ijo.2012.1420 (2012).

36 Li, A. et al. Circadian gene Clock contributes to cell proliferation and migration of glioma and is directly regulated by tumor-suppressive miR-124. FEBS letters 587, 2455–2460, doi:10.1016/j.febslet.2013.06.018 (2013).

37 Flavahan, W. A. et al. Brain tumor initiating cells adapt to restricted nutrition through preferential glucose uptake. Nat Neurosci 16, 1373–1382, doi:10.1038/nn.3510 (2013).

38 Takebe, N. et al. Targeting Notch, Hedgehog, and Wnt pathways in cancer stem cells: clinical update. Nature reviews. Clinical oncology 12, 445–464, doi:10.1038/nrclinonc.2015.61 (2015).

39 Seoane, J., Le, H. V., Shen, L., Anderson, S. A. & Massague, J. Integration of Smad and forkhead pathways in the control of neuroepithelial and glioblastoma cell proliferation. Cell 117, 211–223 (2004).

40 Neuzillet, C. et al. Targeting the TGFbeta pathway for cancer therapy. Pharmacol Ther 147, 22–31, doi:10.1016/j.pharmthera.2014.11.001 (2015).

41 Huang, P. H. et al. Quantitative analysis of EGFRvIII cellular signaling networks reveals a combinatorial therapeutic strategy for glioblastoma. Proceedings of the National Academy of Sciences of the United States of America 104, 12867–12872, doi:10.1073/pnas.0705158104 (2007).

42 Inda, M. M. et al. Tumor heterogeneity is an active process maintained by a mutant EGFR-induced cytokine circuit in glioblastoma. Genes & development 24, 1731–1745, doi:10.1101/gad.1890510 (2010).

43 Shinojima, N. et al. Prognostic value of epidermal growth factor receptor in patients with glioblastoma multiforme. Cancer research 63, 6962–6970 (2003).

44 Padfield, E., Ellis, H. P. & Kurian, K. M. Current Therapeutic Advances Targeting EGFR and EGFRvIII in Glioblastoma. Front Oncol 5, 5, doi:10.3389/fonc.2015.00005 (2015).

45 Morrissey, K. M., Yuraszeck, T., Li, C. C., Zhang, Y. & Kasichayanula, S. Immunotherapy and Novel Combinations in Oncology-Current Landscape, Challenges and Opportunities. Clin Transl Sci, doi:10.1111/cts.12391 (2016).

46 Huntley, R. P. et al. The GOA database: gene Ontology annotation updates for 2015. Nucleic Acids Res 43, D1057–1063, doi:10.1093/nar/gku1113 (2015).

47 Pruitt, K. D. et al. The consensus coding sequence (CCDS) project: Identifying a common protein-coding gene set for the human and mouse genomes. Genome research 19, 1316–1323, doi:10.1101/gr.080531.108 (2009).

48 Gonzalez-Perez, A. et al. IntOGen-mutations identifies cancer drivers across tumor types. Nat Methods 10, 1081–1082, doi:10.1038/nmeth.2642 (2013).

49 Cline, M. S. et al. Integration of biological networks and gene expression data using Cytoscape. Nature protocols 2, 2366–2382, doi:10.1038/nprot.2007.324 (2007).

